# Calcium imaging reveals coordinated simple spike pauses in populations of Cerebellar Purkinje cells

**DOI:** 10.1101/051730

**Authors:** Jorge E. Ramirez, Brandon M. Stell

**Author notes:** To whom correspondence should be addressed: Brain Physiology Lab - Université Paris Descartes, 45, rue des Saints Péres, 75270 Paris Cedex 06, France — (Brandon M. Stell).

## Abstract

The brain’s control of movement is thought to involve coordinated activity between cerebellar Purkinje cells. The results reported here demonstrate that somatic Ca^2+^ imaging is a faithful reporter of Na^+^-dependent “simple spike” pauses and enables us to optically record changes in firing rates in populations of Purkinje cells. This simultaneous calcium imaging of populations of Purkinje cells reveals a striking spatial organization of pauses in Purkinje cell activity between neighboring cells. The source of this organization is shown to be the presynaptic GABAergic network and blocking GABA_*A*_Rs abolishes the synchrony. These data suggest that presynaptic interneurons synchronize (in)activity between neighboring Purkinje cells and thereby maximize their effect on downstream targets in the deep cerebellar nuclei.

## Introduction

*In vivo* measurements from the molecular layer of the cerebellar cortex show that active parallel fibers directly excite Purkinje cells and, via inhibition from molecular layer interneurons (MLIs), pause spiking in Purkinje cells lateral to active parallel fibers (Andersen et al. 1964; Cohen and Yarom 2000; Dizon and Khodakhah 2011). This dual action of parallel fibers likely explains why sensory stimulation produces both positive and negative correlations between *in vivo* recordings of activity in Purkinje cells and their targets in the deep cerebellar nuclei (DCN) (Person and Raman 2012b). Lateral inhibition could also explain the spontaneous pauses in Purkinje cell spiking, lasting longer than several hundred milliseconds, that are routinely recorded *in vivo* under a variety of different recording conditions in several different species (Bell and Grimm 1969; Crepel 1972; Loewenstein et al. 2005; Schonewille et al. 2006; Bosman et al. 2010; Kitamura and Häusser 2011).

Coordinated activity in Purkinje cells has been shown to have a large impact on the output of the DCN *in vivo* (Person and Raman 2012a; Chaumont et al. 2013). Using electrophysiological techniques, several groups have recorded highly correlated individual action potentials in Purkinje cells from a variety of species *in vivo* (Bell and Grimm 1969; Ebner and Bloedel 1981; de Solages et al. 2008; Person and Raman 2012b), but resolving this coordination has been limited by electrophysiology techniques and therefore the spatial organization of coordinated pauses have not been described previously. Moreover, despite somatic expression of voltage gated Ca^2+^ channels in Purkinje cells (Callewaert et al. 1996), monitoring individual action potentials with Ca^2+^ imaging has previously proven difficult (Lev-Ram et al. 1992; Miyakawa et al. 1992). However, Purkinje cells fire trains of action potentials and it has been suggested that it may be possible to detect the transitions between periods of spiking and silence (Lev-Ram et al. 1992; Miyakawa et al. 1992) and that those transitions could potentially be used to monitor pauses in Purkinje cell spiking as previously shown possible in other smaller neurons (Franconville et al. 2011).

Several groups have recently shown that when pauses in simple spike activity are imposed on populations of Purkinje cells *in vivo*, they can produce robust motor output (Lee et al. 2015; Heiney et al. 2014). In addition, there is evidence that pauses in Purkinje cell spiking are involved in cerebellar dependent learning paradigms (Lee et al. 2015; Jirenhed et al. 2007; Jirenhed and Hesslow 2011; Johansson et al. 2015; Johansson et al. 2014). Therefore, the ability to monitor these pauses in populations of Purkinje cells could open new possibilities for understanding their role in cerebellar processing.

## Results

We recorded from Purkinje cells in cerebellar slices using the whole-cell patch clamp technique and loaded the cells for ≈ 20 minutes with 50-100 *μ*M Oregon Green BAPTA-1 (OGB1) in the recording pipette. After retracting the whole-cell electrode we lowered another electrode onto the cell and performed loose cell-attached recordings of the spontaneous action potentials while simultaneously imaging fluorescence with a CCD camera (Materials and Methods). Small, spontaneous changes in fluorescence (4.3 ± 0.9% ΔF/F_0_ from peak to peak; n = 12 cells), were observed over several minutes and pauses in Purkinje cell activity, recorded with the cell-attached electrode, coincided with sharp decays in fluorescence (Fig. 1). We found that the pauses in spiking could be readily detected off-line using only the fluorescence signal by plotting the second time derivative of the calcium signals and using a simple threshold (Fig. 1B, C). As shown in figure 1, these threshold crossings can be plotted versus time to construct an idealized binary diagram that reports when the Purkinje cell is firing (shaded regions in figure 1 C) or has paused its firing (unshaded regions in figure 1 C) and they appeared to correspond remarkably well to the electrophysiological records of the simple-spike activity.

**Figure 1:**
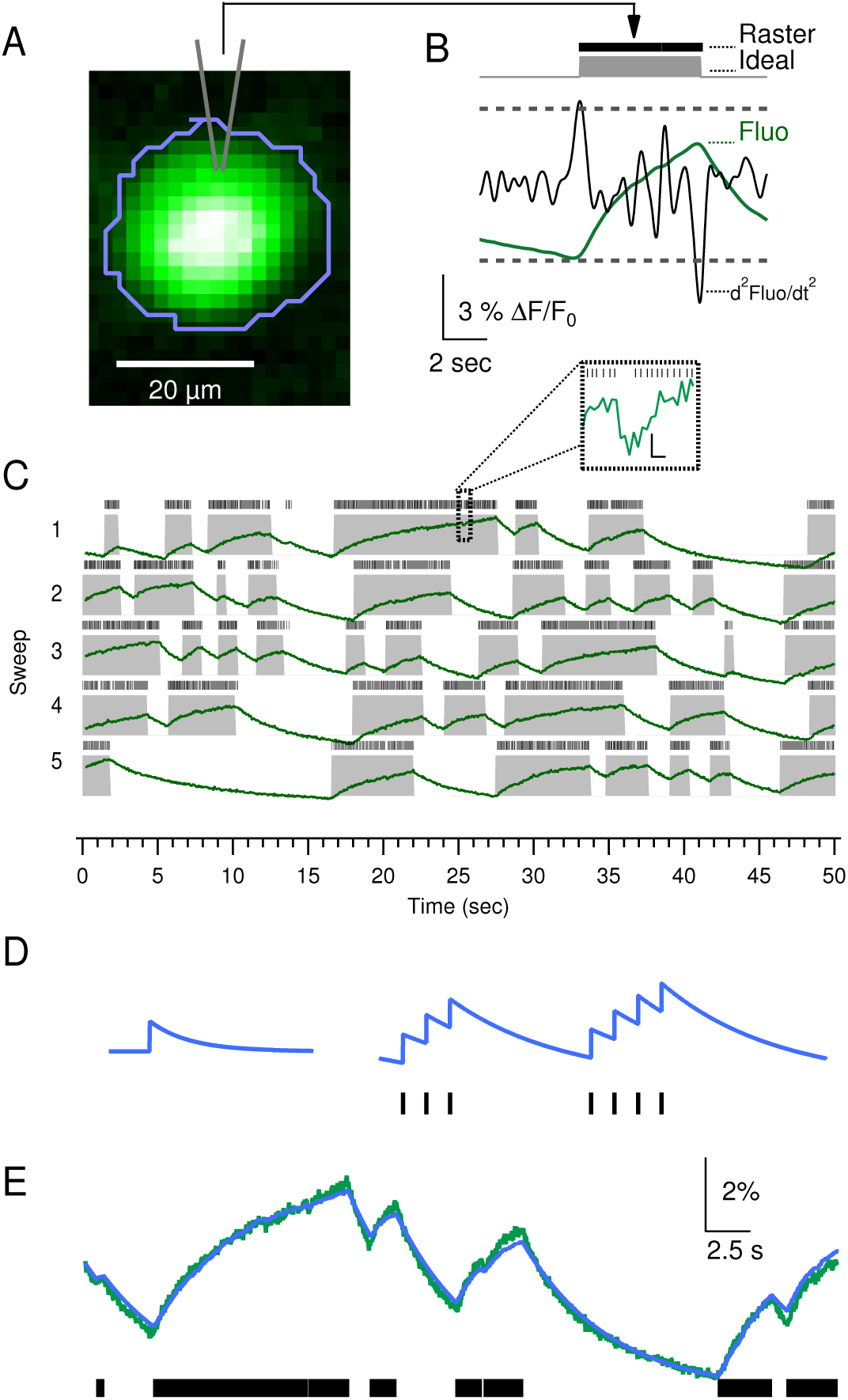
Somatic calcium imaging reports Purkinje cell simple spike activity. **A** Purkinje cell preloaded with 100 *μ*M OGB1. Image scaled to brightest and dimmest pixels (brightest = 2200 counts; max counts = 14 bits). **B** Raster plot of spontaneous activity from a cell-attached electrode on the same cell shown in A with corresponding smoothed (binomial smoothing, ten operations) calcium signal (green), its second derivative (black), and the activity diagram (gray) constructed from threshold (dashed lines) crossings. **C A** longer timescale of the same data shown in B. The cell was firing with a median firing frequency of 20.3 Hz and the inset calls out section of the recording with an inter-spike interval of 151 ms (≈ 3 missing APs). Scale bars of inset indicate 0.3% ΔF/F_0_ and 100 ms. **D** Schematic of an idealized calcium response with a single exponential decay on the left and the addition of seven such responses at the times indicated by the points in the raster plot below in black. **E** Green trace shows a portion of the data from the calcium recording depicted in the first sweep of panel C and the corresponding raster plot of the action potentials recorded from the cell in cell-attached mode in black below. Blue trace shows the sum of a single exponential with a tau of 4.9 s and an amplitude of 0.06% for each of the action potentials in the raster plot.

Given that the somatic calcium appeared to correlate extremely well to the pauses in simple spike activity, we attempted to determine whether the calcium signal could also be used to monitor spiking rate during the active periods. To do this we recreated the calcium signal using only the timing of the action potentials recorded in cell-attached mode. We used a kernel with an instantaneous rise and single exponential decay (Fig. 1D, left) and convolved it with the spike train that was recorded in the cell-attached recordings (Fig. 1D, right). We then used a least squares fitting routine to adjust the asymptote, decay time constant, and amplitude of the kernel calcium response until we found the best fit to the raw calcium recording as can be seen in Fig. 1E. The convolution of the kernel calcium transient with the simultaneously recorded action potentials resulted in remarkably accurate fits of the raw calcium responses and allowed us to estimate the average calcium response of each cell to a single simple spike. This analysis reported a mean simple-spike induced ΔF/F_0_ of 0.1 ± 0.03 % with a mean decay time of 7.2 ± 2.1 s (n = 6 cells).

The calcium transients produced by single action potentials were surprisingly similar between cells and encouraged us to further investigate whether calcium recordings alone could be used to determine changes in simple spike firing rate in Purkinje cells. To do this we focused on a simple change in firing rate and analyzed all transitions from silent to spiking periods in 7 cells. The ΔF/F_0_ of the calcium recordings for these transitions was calculated by fitting a line to the last ten points before the transition (Fig. 2A) and then subtracting that line from the fluorescence trace (Fig. 2B). The slope of the ΔF/F_0_ after the transition was plotted against the average firing rate recorded with the cell-attached electrode for each transition for all 7 cells. As can be seen in Fig. 2C, the firing rate recorded in the cell-attached electrode correlated strongly with change in fluorescence recorded by the calcium imaging. Remarkably, this correlation was obvious both for individual cells as well as between cells, indicating that somatic calcium alone is an excellent reporter of simple-spike firing rate changes. Furthermore these data indicate that individual simple-spikes trigger similar magnitudes of Ca^2+^ entry into the soma of different Purkinje cells.

**Figure 2:**
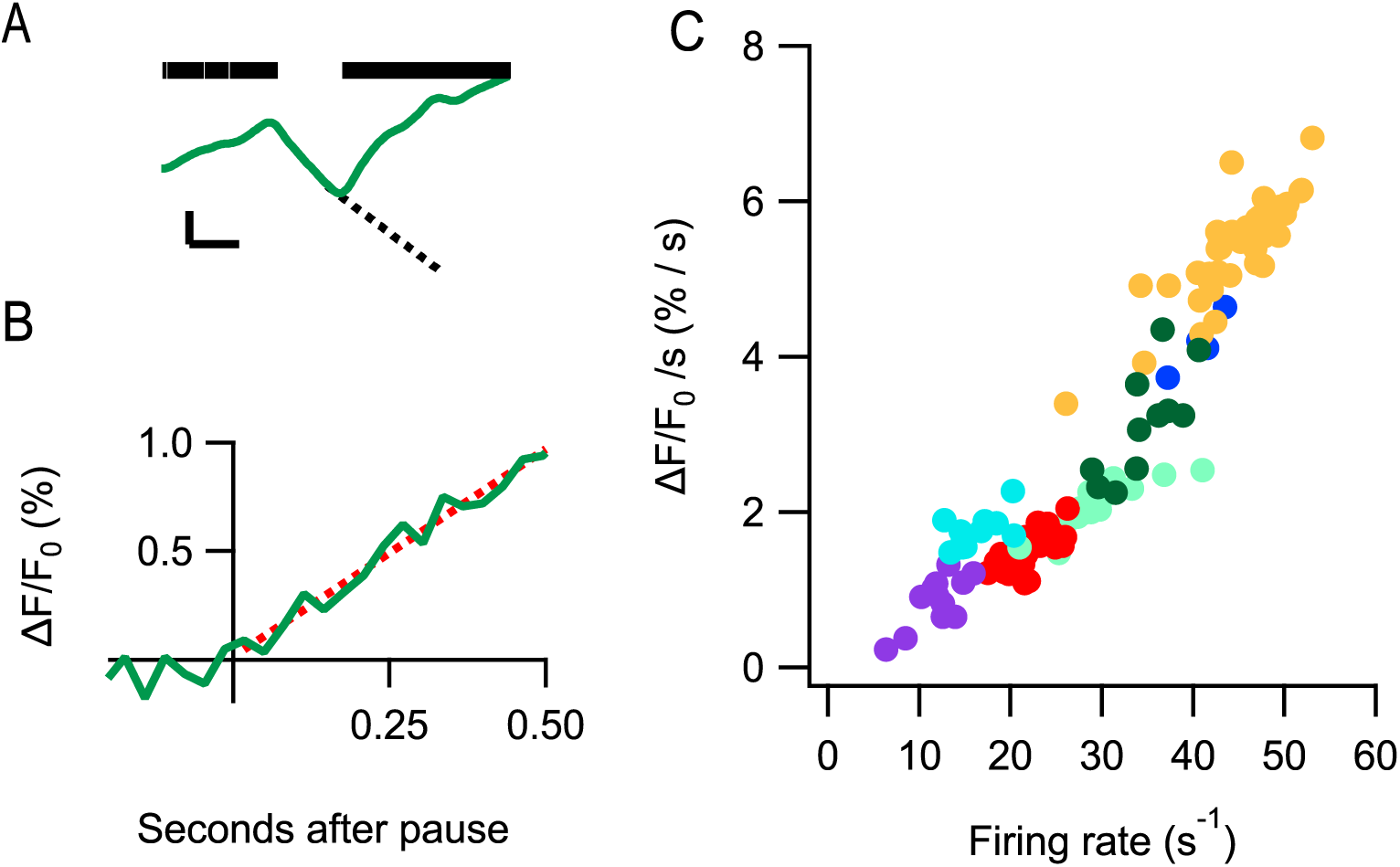
Somatic calcium imaging reports Purkinje cell firing frequency. **A** Average raw fluorescence recorded from the soma of a Purkinje cell (green) and a raster plot of the corresponding action potentials recorded in cell-attached mode (black). Dashed line is a linear fit to the average fluorescence recorded from the soma during the ten frames just prior to the end of the pause in spiking. Scale bar = 10 counts and 1 sec. **B** Same data from A converted to ΔF/F_0_ using the dashed line in A as F_0_. Dashed red line is a linear fit to the data recorded in the 500 msec following the pause. **C** Data from seven cells (indicated by different colors; red points are from the cell shown in A and B) in which the change in ΔF/F_0_ during the 500 msec following a pause (slope of dashed red line in B) is plotted against the average firing rate ( 1 / average inter-spike interval) of action potentials recorded in cell-attached mode over the same period.

A potential problem with using somatic calcium as a reporter of simple spike activity in Purkinje cells is that these cells also receive a climbing fiber input that produces large calcium transients. We therefore measured the relative effect of simple vs complex spikes on somatic calcium. In Fig. 3B we show simultaneous calcium and cell-attached measurements recorded while we stimulate a climbing fiber input onto a Purkinje cell. As can be seen in the individual sweeps shown in Fig. 3B, and consistent with previous reports (Eilers et al. 1995; Lev-Ram et al. 1992; Kano et al. 1992; Tank et al. 1988), the climbing fiber activation produces a large change in fluorescence in the Purkinje cell dendrites and a much smaller signal in the soma (ΔF/F_0_ 36.2 ± 5.4% in dendrites and 1.9 ± 0.2 % in soma; n = 12 cells). This effect can be seen more clearly in the right panel of Fig. 3C, which depicts the average peak response of a single Purkinje cell to 15 climbing fiber stimuli (one every 20 seconds). In this image, it is clear that the large climbing fiber response recorded in the dendrites is not recorded in the soma. By contrast the same analysis of the simple spike calcium response (Fig. 3C left) shows a large response in the soma that is much smaller in the dendrites. Therefore, as shown in (Fig. 3D), climbing fiber calcium responses can be easily distinguished from calcium responses induced by trains of simple-spike activity based on the ratio of dendritic to somatic fluorescence.

**Figure 3:**
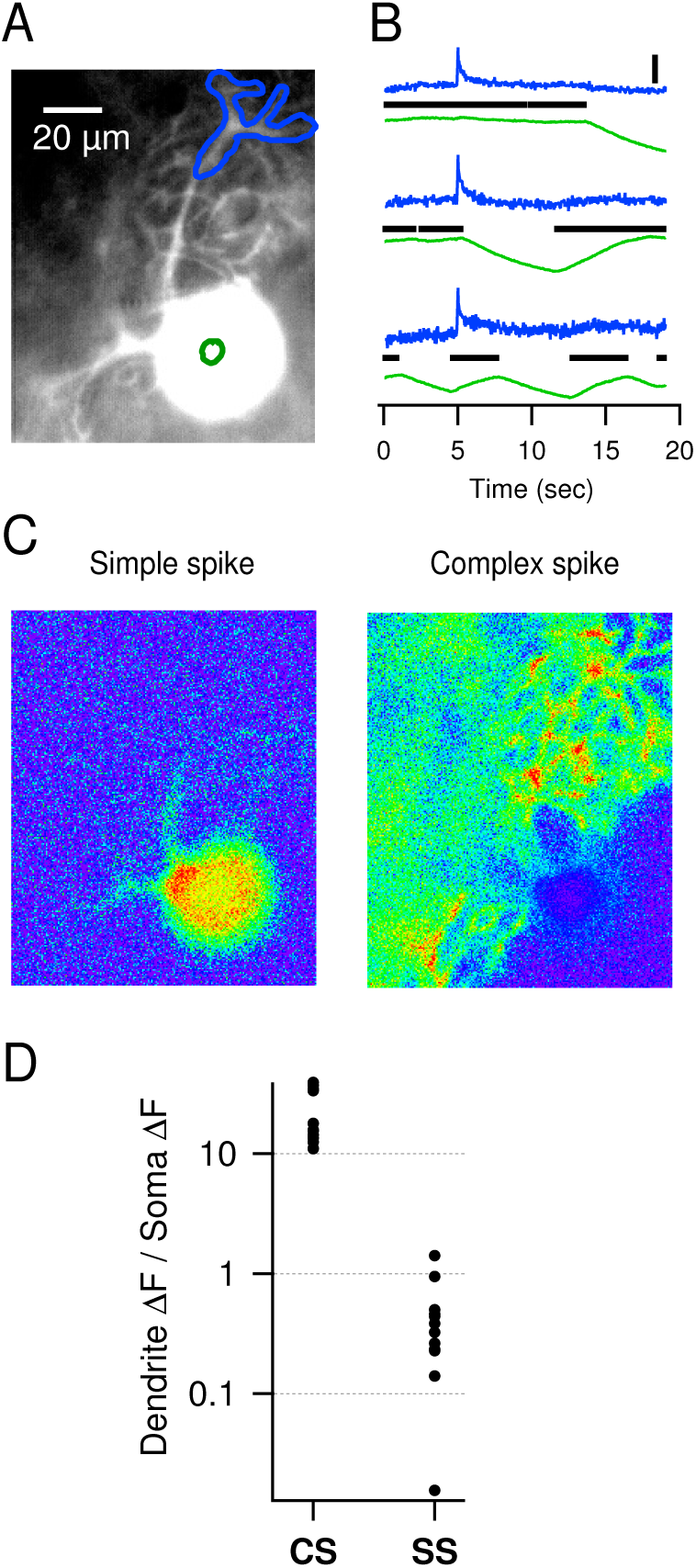
Somatic calcium reports simple spikes but not complex spikes. **A** Purkinje cell loaded with 100 *μ*M OGB-1 via the patch pipette for 20 minutes. **B** ΔF/F_0_ of fluorescence recorded from the two regions of interest shown in A (dendrites in blue and soma in green) and the corresponding raster plot of action potentials recorded in cell-attached mode (black). Climbing fiber stimulation (at 5 sec) produces a large increase in fluorescence in the dendrites that is not recorded in the soma. Scale bar = 20 % ΔF/F_0_. **C** Peak response of cell shown in **A** and **B** to simple or complex spikes. Images display the peak in the average ΔF/F_0_ of 14 repetitions for the simple spikes and 28 repetitions for the complex spikes. F_0_ is the average for each pixel taken across the five frames just prior to either simple spike train or the complex spike. Same scale bar as in A. Each image is scaled to show min to max fluorescence from 0 - 10% ΔF/F_0_ **D** Ratio of the change in dendritic over somatic fluorescence for complex and simple spikes plotted on a logarithmic y-axis. With a ratio >10, complex spikes can be easily distinguished from simple spikes.

Since somatic calcium was determined to be an excellent reporter of simple spike activity, we used it to monitor correlations in activity between multiple Purkinje cells simultaneously (Fig. 5A). Correlations between two cells could be measured by comparing the average Pearson’s correlation indexes from cross-correlations performed on simultaneous Ca^2+^ recordings to the cross-correlations performed on Ca^2+^ recordings from the same cells at different times (see Materials and Methods and Fig. 4). By performing this analysis on the second time derivative of the calcium recordings (shown above to be an indicator of transitions into and out of pauses in Purkinje cell spiking; Fig. 1B and C), correlations among transitions were revealed between neighboring cells in sagittal planes of either transverse or sagittal slices (P < 0.05 for all pairs; n = 10). Interestingly, we didn’t record correlations in pairs of cells that were in different sagittal planes of the same transverse slices (P > 0.05 in 5 of 5 pairs). To determine the spatial extent of correlated Purkinje cell pauses in the sagittal plane, correlations were also performed on data from a second set of electrophysiology experiments (Materials and Methods) in which cell-attached recordings were made from pairs of Purkinje cells. This method allowed us to examine Purkinje cell pairs beyond the field of view of our imaging system (separated by up to 350 um). This method also revealed strong correlations in pauses of activity in neighboring Purkinje cells up to 200 *μ*m apart (Fig. 5B) which is the approximate length of the main branch of MLI axons in the cerebellar cortex (211 *μm;* de San Martin et al. 2015).

**Figure 4:**
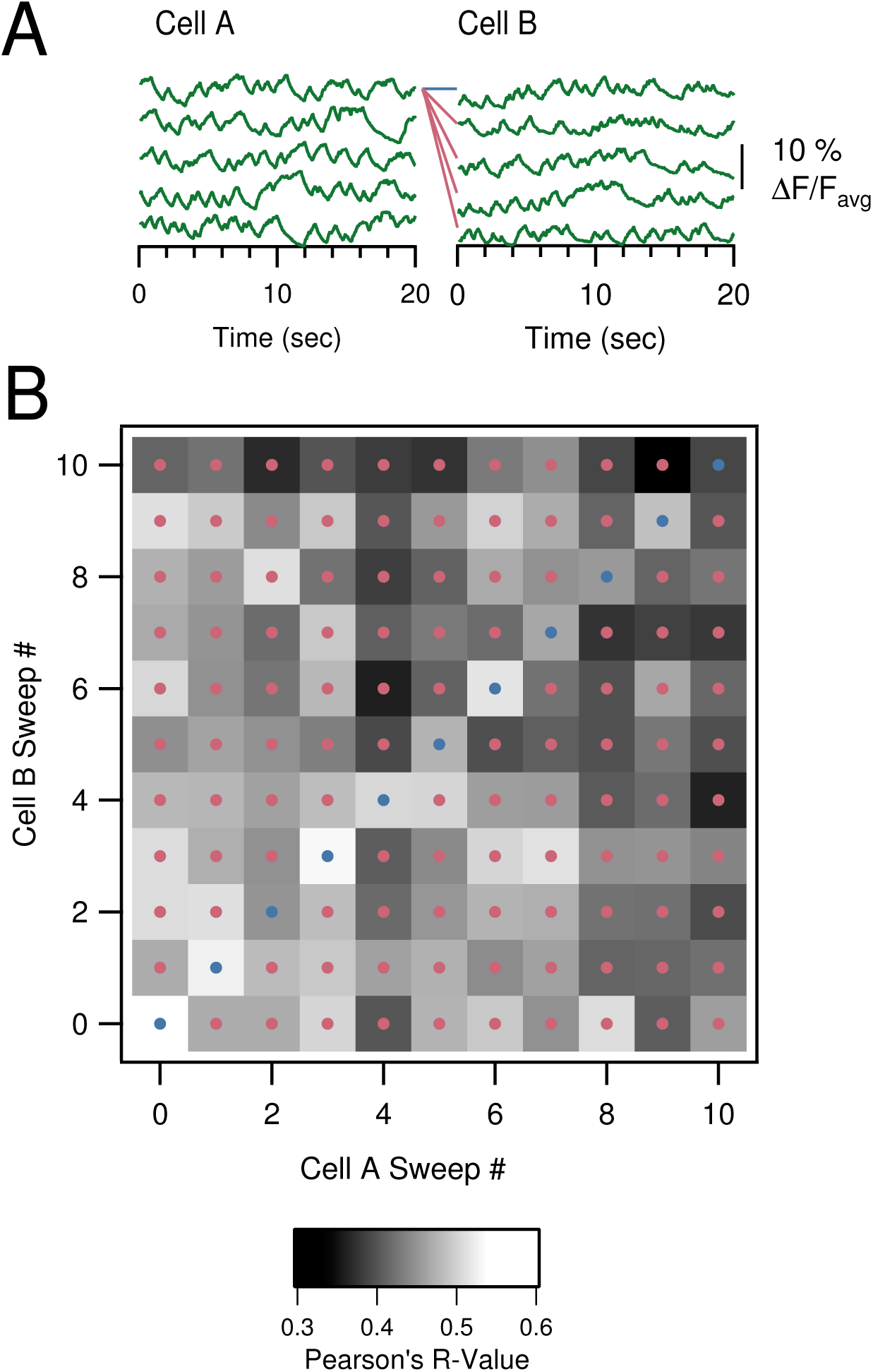
**A** Simultaneous calcium recordings from two cells. Blue line indicates a paired correlation between the first recordings in the 2 cells: point 0,0 in B. Red lines indicate the shuffled correlations for the first sweep in cell A: column 0 in B. **B** Matrix of Pearson’s correlation coefficients calculated for all simultaneous recordings (unity line / blue dots) and shuffled recordings (pink dots). To determine whether there was a correlation between two cells, a Student’s t-test was performed between the resulting populations of simultaneous and shuffled correlation coefficients and a significant difference was interpreted as a significant correlation.

**Figure 5:**
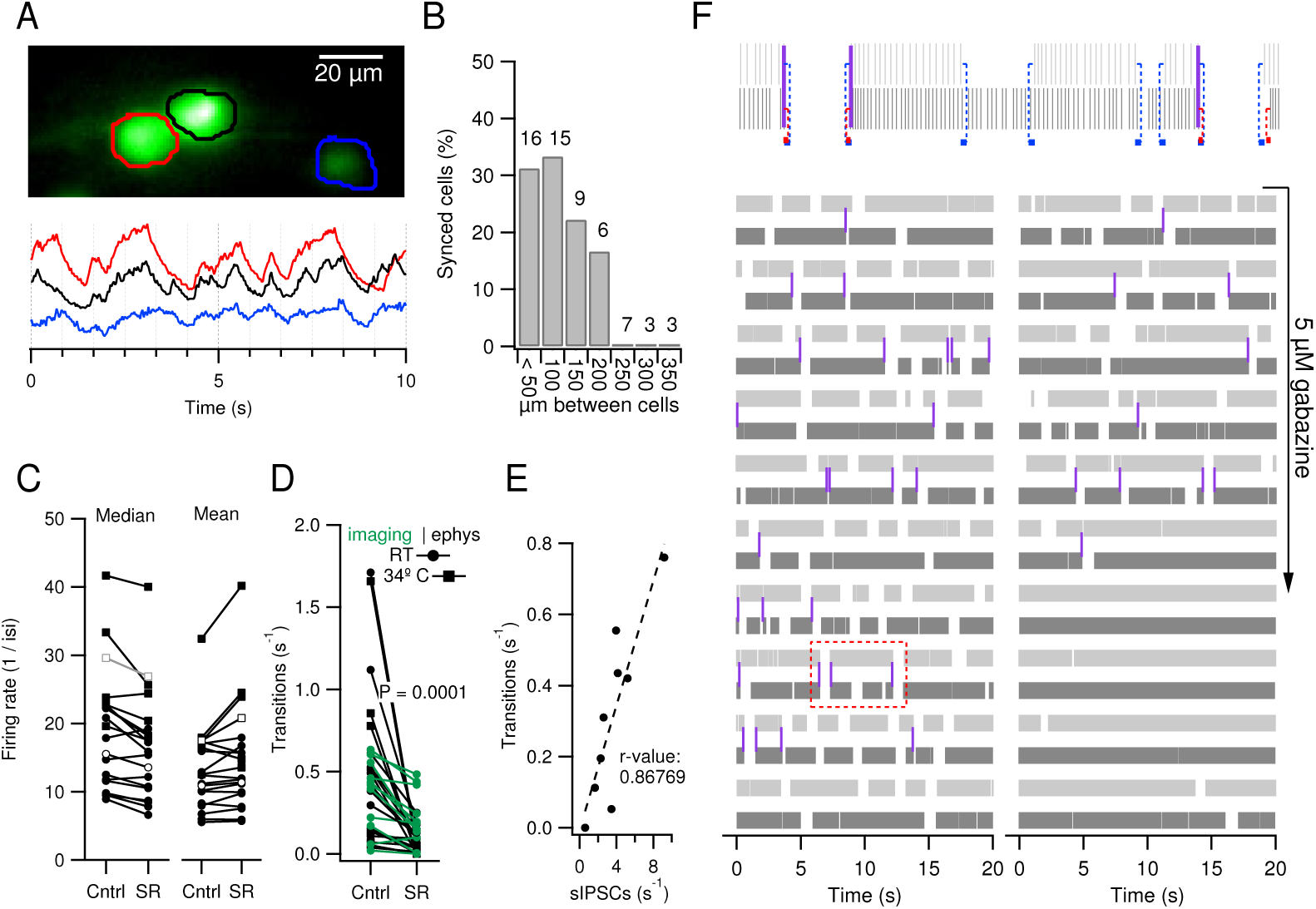
GABA-A receptors correlate activity between direct Purkinje cell neighbors in the same sagittal plane. **A** Three Purkinje cell soma in roughly the same sagittal plane of a transverse slice (Purkinje cell dendrites descending into the plane of the slice) individually preloaded with 100 *μ*M OGB-1 for 5 min each. Lower panel plots ΔF/F_0_ from the ROIs drawn on the upper image. **B** Percentage of pairs of cells recorded in cell-attached mode that show significant synchrony in transitions into and out of pauses (lasting longer than 3 times their median firing frequency) as a function of distance between somas. The number of pairs recorded at each distance is indicated above each bar. **C** Firing rate (median and mean) recorded with cell-attached electrodes in control conditions and after washing 5 *μ*M gabazine into the bath. As in D, boxes represent recordings made at 34°C and circles represent recordings made at room temperature; open symbols represent the averages for each condition. **D** Rate of transitions to and from pauses in control conditions and in the presence of 5 *μ*M gabazine. Gabazine significantly reduced the frequency of transitions (P = 0.0001). Electrophysiology data shown in black and imaging data shown in green were not significantly different in control conditions nor in the presence of gabazine (P >0.05). **E** Transitions to and from pauses lasting longer than 3 times the median inter-spike interval recorded in loose cell-attached recordings plotted versus spontaneous IPSC frequency recorded in subsequent whole-cell recordings from the same cells (n = 9 cells). **F** Raster plot of action potentials detected in two Purkinje cells (each cell in different shades of gray) with loose cell-attached recordings from two neighboring Purkinje cells (represented in different shades of gray. The upper panel illustrates an expanded section of the lower raster (shown in red dash) and demonstrates how simultaneous transitions between pauses were detected: one median inter-spike interval is indicated with a dashed line in blue or red at the beginning and end of a pause (any inter-spike interval >3 times the median) that indicate a simultaneous transition when overlapping (purple bars). Gabazine is washed into the recording chamber where indicated and clearly decreases the frequency of simultaneous transitions.

It has been shown previously that MLIs play a large role in controlling the timing of Purkinje cell simple spikes (Häusser and Clark 1997), and that individual MLIs can transition Purkinje cells in and out of spiking periods (Oldfield et al. 2010). We found that blocking GABA_A_Rs with gabazine (5 *μ*M) caused a large reduction in the frequency of spontaneous transitions into and out of pauses recorded either with Ca^2+^ imaging (0.34 ± 0.06 Hz in control and 0.20 ± 0.05 Hz in gabazine; n = 12; P = 0.001, Paired Student’s t-test) or with loose cell-attached electrodes (0.51 ± 0.05 Hz in control and 0.15 ± 0.04 Hz in gabazine, n = 4; P = 0.004, Paired Student’s t-test; Fig. 5D). The frequency of transitions was similar for loose cell-attached and calcium recordings in both control conditions (0.34 ± 0.06 Hz for cell-attached vs 0.51 ± 0.05 Hz for imaging; P = 0.06; Student’s t-test) and in gabazine (0.20 ± 0.05 vs 0.15 ± 0.04; P = 0.37, Student’s t-test), indicating that calcium imaging does not affect the frequency of transitions. Since this indicates that GABA_A_R activation influences Purkinje cell pausing and it is known that GABAergic synapses are temperature sensitive, we tested whether elevated temperatures affect Purkinje cell pausing and found no difference in frequency when recordings were made at room temperature versus elevated temperature (P = 0.49; n = 5 cells recorded at 34°C and 14 at room temperature). Consistent with previous results (Häusser and Clark 1997) when we examined the instantaneous firing rate in cells recorded with cell-attached electrodes (calculated using the inverse of the inter-spike intervals recorded over several minutes), we found that the mean firing rate was increased by gabazine (Fig. 5C; 13.2 ± 1.5 spikes / sec in control vs 14.6 ± 2.0 spikes / second in gabazine; P = 0.04; paired Student’s t-test; n = 18 cells), but since the mean spike rate is influenced by the number and length of pauses in spiking and we show that gabazine decreases the frequency of pauses, this results is perhaps not surprising. However, since action potentials always greatly outnumber pauses in firing by at least one order of magnitude (Fig. 5C and D), the median spike rate is only subtly affected by pauses and therefore more accurately reflects the spike rate during spiking periods. Surprisingly, we found that gabazine slightly decreased the median firing rate (Fig. 5C; 18.6 ± 2.1 spikes / sec in control vs 16.5 ± 1.9 spikes / second in gabazine; P = 0.001; paired Student’s t-test; n = 18 cells) indicating that tonic activation of GABA_A_Rs acts to increase the firing rate of Purkinje cells. By performing a series of experiments in which the frequency of transitions in a given Purkinje cell was measured with a loose cell-attached electrode before breaking into the cell and recording the frequency of spontaneous IPSCs, we found that overall activity of the presynaptic GABAergic network plays a major role in controlling Purkinje cell pausing because the number of transitions was directly proportional to the frequency of spontaneous IPSCs (Fig. 5E; r-value 0.87; n = 9 cells). Altogether these data suggest that GABA released by interneurons can pause Purkinje cell spiking and that MLIs are participating in controlling the correlations in pauses. Therefore, to directly test the involvement of MLIs in correlating Purkinje cell pausing, we washed gabazine onto pairs of Purkinje cells simultaneously recorded in the loose cell-attached configuration (Fig. 5F) and found that correlations in Purkinje cell pauses were always abolished by the GABA_A_R blocker gabazine (Fig. 5F; P < 0.05 in control and P > 0.05 after gabazine for all pairs; n = 7; see Materials and Methods).

## Discussion

The results presented here demonstrate that Ca^2+^-imaging can be used to monitor simple spike activity in Purkinje cells and opens new options to explore the morphological organization of Purkinje cell (in)activity *in vivo*. The technique also reveals that simple spikes in Purkinje cells produce a surprisingly consistent rise in somatic calcium concentration (even between Purkinje cells) and therefore the technique can be used to examine pauses in Purkinje cell simple spike trains as well as changes in firing rate as shown in figure 2C.

Using this technique in slices of the cerebellar cortex we reveal a coordination of Purkinje cell pauses that appears to be structured by the presynaptic GABAergic network to be perpendicular to the parallel fiber axis. Pauses in Purkinje cell simple-spiking have been recorded before both in the slice and *in vivo* (Bell and Grimm 1969; Crepel 1972; Loewenstein et al. 2005; Schonewille et al. 2006; Yartsev et al. 2009; Bosman et al. 2010; Oldfield et al. 2010; Kitamura and Häusser 2011) but coordination between pauses in spontaneous simple spikes of Purkinje cells has not been previously reported. The output neurons in the DCN respond strongly to coordinated inactivity of Purkinje cells (Aizenman and Linden 1999; Heiney et al. 2014; Person and Raman 2012a) and therefore, unlike coordinated small variabilities in Purkinje cell spiking (De Schutter and Steuber 2009), the coordination of several missing action potentials in Purkinje cells that we report here is likely to have a large impact on the output of information from the cerebellum.

It has been shown previously in the rodent that the convergence of Purkinje cells onto the DCN varies between 20 - 50 Purkinje cells per DCN cell (Person and Raman 2012a), and although the magnitude of the resulting inhibition is modulated by the exact timing of individual simple spikes of Purkinje cells, it is clear that regardless of the timing of their spikes the net effect of Purkinje cells is to produce constant inhibition of the spontaneous activity of the DCN (Person and Raman 2012a). Therefore pauses in Purkinje cell activity coordinated between sets of Purkinje cells synapsing onto the same DCN cell would produce a strong signal at that level. It is shown here that sets of Purkinje cells located in the same sagittal plane show coordinated pauses in activity and is consistent with previous reports suggesting that Purkinje cell terminals originating from Purkinje cells located in the same sagittal planes likely synapse onto many of the same cells in the DCN (Sugihara et al. 2009).

## Materials and Methods

### Slice Preparation

Twelve-to sixteen-day old Sprague-Dawley rats were anesthetized with isoflurane and decapitated before the cerebellum was removed. Sagittal or transverse slices of the vermis, 200 *μ*m thick, were cut using a Leica VT 1000S vibratome (Nussloch) in ice-cold artificial cerebrospinal fluid (ACSF) bubbled with 95% oxygen and 5% carbon dioxide. ACSF contained the following (in mM): 130 NaCl, 2.4 KCl, 1.3 NaH_2_PO_4_, 1 MgCl_2_, 2 CaCl_2_, 10 glucose, 26 NaHCO_3_ and 1-3 kynurenic acid. Slices were incubated for 1 h at 34°C in ACSF (without the kynurenic acid) and then kept at room temperature until transferred to the electrophysiological recording chamber.

### Electrophysiological Recordings

The recording chamber was continuously perfused at a rate of 1–2 mL/min with the same ACSF (and in the presence of 10 *μ*M gabazine when indicated below). Recordings were made at room temperature. The Purkinje cell layer and the molecular layer of the cerebellar cortex were visualized using an Axioskop microscope (Zeiss) equipped with a 63X/0.9 water immersion objective. Signals were recorded with an EPC-10 amplifier (HEKA Elektronik) with Patchmaster software v2.32. All cell-attached recordings were made in voltage clamp mode with ACSF in the pipette and 0 mV holding potential. Internal solution was prepared form a stock solution containing (in mM) 165 K-gluconate, 2.4 MgCl_2_, 10 HEPES, 1 EGTA, 0.1 CaCl_2_, 2 Na_2_-ATP, 0.4 Na-GTP pH to 7.4 with KOH, 327 mOsm/KgH_2_O and 50 - 100 *μ*M OGB1 was added (diluting the solution by 10%). For whole-cell recordings in which imaging was not performed this solution was diluted by 10% with distilled water. Borosilicate glass pipettes were pulled using a HEKA Elektronik pipette puller and had an open tip resistance of ~3 MΩ for cell-attached and whole-cell recordings.

### Ca^2+^ Imaging

Purkinje cells were loaded with 50 - 100 *μ*M OGB1 hexapotassium salt added to the pipette solution for ~20 minutes with a whole-cell recording on the soma in voltage clamp mode before pipettes were retracted to form an outside-out patch. The formation of an outside-out patch on the pipette indicated that the membrane resealed on the pipette and therefor also on the soma, thereby trapping the indicator in the cell. Cells were illuminated with a 470 nm LED controlled with a Cairn Optoled system (Cairn Research Ltd). Light from the LED was sent down the epifluorescence pathway and reflected off of a T495LP dichroic mirror (Chroma Technology Corp, USA) onto the preparation. Fluorescence light passed through the dichroic mirror and was filtered with an emission filter (ET525/50M; Chroma Technology Corp, USA) and detected with an ANDOR iXON CCD camera (model 887) at a frame-rate of 31.5 Hz with 30 ms exposures. Changes in average fluorescence were analyzed off-line in Igor Pro (Wavemetrics) with ROIs hand-drawn around the soma using custom functions. ΔF/F_0_ for pauses in simple spiking were calculated by fitting a single exponential to the decay at the beginning of pauses and the asymptote of the fitted exponential was used as F_0_.

### Correlation analysis of imaging data

20 second long acquisitions were analyzed for each correlation. Average fluorescence within a region of interest (ROI) drawn surrounding each soma was computed individually for each ROI in each frame. The average fluorescence from each ROI was plotted vs time and smoothed using a binomial smoothing (10 - 50 operations depending on noise in the signal; see the “smooth” function in Igor Pro version 6.1). The first and second derivatives were computed for the smoothed data. The first derivative reports the activity state of the cell (whether or not it is firing action potentials) whereas the second derivative reports the changes in Purkinje cell activity between active and silent states. Pearson’s correlation coefficients from cross-correlations were computed (using the built-in “statscorrleation” function in Igor Pro version 6.1) between the second derivatives of calcium recordings collected simultaneously from different cells (Fig. 4A) and resulted in simultaneous coefficients for each series (y = x and blue dots in Fig. 4B). Pearson’s correlation coefficients were also computed between each sweep and those from the other cell that were not recorded simultaneously, resulting in shuffled correlations (y ≠ x pink dots in Fig. 4A). A student’s t-test was then performed between the simultaneous coefficients and the shuffled coefficients to determine significance of any correlation between two simultaneously recorded cells.

### Correlation analysis of electrophysiology data

Raster plots were created by detecting threshold crossings using Neuromatic in Igor Pro. Inter-spike intervals (ISIs) were calculated by differentiating the spike timings of the rasters. We considered the median as the parameter which best reflected the central tendency of the ISI distributions and used it to describe the most regular spiking behavior of the cells. Then, we defined pauses as quiescent times which lasted more than 3 times the median (in other words, missing at least two regular action potentials), in order to avoid detecting irregular spiking patterns caused by jittered or misfired spikes. Windows of one median ISI were then attached to the beginning and end of pauses in individual recordings (Fig. 5F upper inset) and when these windows overlapped in simultaneous recordings they were considered to be synchrounous transitions (purple in Fig. 5F). We then randomly reordered the trains of recorded ISIs 500 times for each recording and performed the same analysis to find the simultaneous transitions when the ISIs occurred randomly. This randomization process produced a normal distribution of overlapping transitions to which we compared the number of simultaneous transitions that were recorded in the experiments. This permutation test allowed us to evaluate whether the coordinated pauses in Purkinje cell spiking were significantly different than random coincidences by performing a a one-sided Z-score test (α = 0.001).

## Acknowledgements

This work was funded by the CNRS. We would like to thank the members of the Brain Physiology Lab (CNRS unit UMR8118) for their comments on the manuscript and/or helpful discussions throughout the project, in particular Michael Graupner who translated some of our fitting routines from Igor into Python. We would also like to thank Boris Barbour for suggesting the method used for determining significance of correlations between Purkinje cells, for comments on the manuscript, and discussions throughout the project.

## References

1. Aizenman, C. D. and D. J. Linden (1999). Regulation of the rebound depolarization and spontaneous firing patterns of deep nuclear neurons in slices of rat cerebellum. Journal of Neurophysiology 82(4), 1697–1709.

2. Andersen, P., J. C. Eccles, and P. E. Voorhoeve (1964). Postsynaptic inhibition of cerebellar Purkinje cells. Journal of neurophysiology 27, 1138–1153.

3. Bell, C. C. and R. J. Grimm (1969). Discharge properties of Purkinje cells recorded on single and double microelectrodes. Journal of Neurophysiology 32(6), 1044–1055.

4. Bosman, L. W. J., S. K. E. Koekkoek, J. Shapiro, B. F. M. Rijken, F. Zandstra, B. van der Ende, C. B. Owens, J.-W. Potters, J. R. de Gruijl, T. J. H. Ruigrok, and C. I. De Zeeuw (2010). Encoding of whisker input by cerebellar Purkinje cells. The Journal of physiology 588(Pt 19), 3757–3783.

5. Callewaert, G., J. Eilers, and A. Konnerth (1996). Axonal calcium entry during fast ‘sodium’ action potentials in rat cerebellar Purkinje neurones. The Journal of Physiology 495 (Pt 3), 641–647.

6. Chaumont, J., N. Guyon, A. M. Valera, G. P. Dugué, D. Popa, P. Marcaggi, V. Gautheron, S. Reibel-Foisset, S. Dieudonné, A. Stephan, M. Barrot, J.-C. Cassel, J.-L. Dupont, F. Doussau, B. Poulain, F. Selimi, C. Léna, and P. Isope (2013). Clusters of cerebellar Purkinje cells control their afferent climbing fiber discharge. Proceedings of the National Academy of Sciences 110(40), 16223–16228.

7. Cohen, D. and Y. Yarom (2000). Cerebellar on-beam and lateral inhibition: two functionally distinct circuits. Journal of neurophysiology 83(4), 19321–1940.

8. Crepel, F. (1972). Maturation of the Cerebellar Purkinje Cells I. Postnatal Evolution of the Purkinje Cell Spontaneous Firing in the Rat. Experimental Brain Research 14, 463–471.

9. de San Martin, J. Z., A. Jalil, and F. F. Trigo (2015). Impact of single-site axonal GABAergic synaptic events on cerebellar interneuron activity. The Journal of General Physiology 146 (6), 477–493.

10. De Schutter, E. and V. Steuber (2009). Patterns and pauses in Purkinje cell simple spike trains: experiments, modeling and theory. Neuroscience 162(3), 816–826.

11. de Solages, C., G. Szapiro, N. Brunel, V. Hakim, P. Isope, P. Buisseret, C. Rousseau, B. Barbour, and C. Léna (2008). High-frequency organization and synchrony of activity in the Purkinje cell layer of the cerebellum. Neuron 58(5), 775–788.

12. Dizon, M. J. and K. Khodakhah (2011). The Role of Interneurons in Shaping Purkinje Cell Responses in the Cerebellar Cortex. The Journal of Neuroscience 31(29), 10463–10473.

13. Ebner, T. J. and J. R. Bloedel (1981). Correlation between activity of Purkinje cells and its modification by natural peripheral stimuli. Journal of Neurophysiology 45(5), 948–961.

14. Eilers, J., G. Callewaert, C. Armstrong, and A. Konnerth (1995). Calcium signaling in a narrow somatic submembrane shell during synaptic activity in cerebellar purkinje neurons. Proceedings of the National Academy of Sciences 92(22), 10272–10276. PMID: 174797661.

15. Franconville, R., G. Revet, G. Astorga, B. Schwaller, and I. Llano (2011). Somatic calcium level reports integrated spiking activity of cerebellar interneurons in vitro and in vivo. Journal of Neurophysiology 106(4), 1793–1805.

16. Häusser, M. and B. A. Clark (1997). Tonic synaptic inhibition modulates neuronal output pattern and spatiotemporal synaptic integration. Neuron 19(3), 665–678.

17. Heiney, S. A., J. Kim, G. J. Augustine, and J. F. Medina (2014). Precise Control of Movement Kinematics by Optogenetic Inhibition of Purkinje Cell Activity. The Journal of Neuroscience 34(6), 2321–2330.

18. Jirenhed, D.-A., F. Bengtsson, and G. Hesslow (2007). Acquisition, Extinction, and Reacquisition of a Cerebellar Cortical Memory Trace. The Journal of Neuroscience 27(10), 2493–2502.

19. Jirenhed, D.-A. and G. Hesslow (2011). Time Course of Classically Conditioned Purkinje Cell Response Is Determined by Initial Part of Conditioned Stimulus. The Journal of Neuroscience 31(25), 9070–9074.

20. Johansson, F., H. A. E. Carlsson, A. Rasmussen, C. H. Yeo, and G. Hesslow (2015). Activation of a Temporal Memory in Purkinje Cells by the mGluR7 Receptor. Cell Reports 13(9), 1741–1746.

21. Johansson, F., D.-A. Jirenhed, A. Rasmussen, R. Zucca, and G. Hesslow (2014). Memory trace and timing mechanism localized to cerebellar Purkinje cells. Proceedings of the National Academy of Sciences 111(41), 14930–14934.

22. Kano, M., U. Rexhausen, J. Dreessen, and A. Konnerth (1992, Apr). Synaptic excitation produces a long-lasting rebound potentiation of inhibitory synaptic signals in cerebellar purkinje cells. Nature 356(6370), 601–604.

23. Kitamura, K. and M. Häusser (2011). Dendritic Calcium Signaling Triggered by Spontaneous and Sensory-Evoked Climbing Fiber Input to Cerebellar Purkinje Cells In Vivo. The Journal of Neuroscience 31(30), 10847–10858.

24. Lee, K., P. Mathews, A. B. Reeves, K. Choe, S. Jami, R. Serrano, and T. Otis (2015). Circuit Mechanisms Underlying Motor Memory Formation in the Cerebellum. Neuron 86(2), 529–540.

25. Lev-Ram, V., H. Miyakawa, N. Lasser-Ross, and W. N. Ross (1992). Calcium transients in cerebellar Purkinje neurons evoked by intracellular stimulation. Journal of Neurophysiology 68(4), 1167–1177.

26. Loewenstein, Y., S. Mahon, P. Chadderton, K. Kitamura, H. Sompolinsky, Y. Yarom, and M. Häusser (2005). Bistability of cerebellar Purkinje cells modulated by sensory stimulation. Nature Neuroscience 8(2), 202–211.

27. Miyakawa, H., V. Lev-Ram, N. Lasser-Ross, and W. N. Ross (1992). Calcium transients evoked by climbing fiber and parallel fiber synaptic inputs in guinea pig cerebellar Purkinje neurons. Journal of Neurophysiology 68(4), 1178–1189.

28. Oldfield, C. S., A. Marty, and B. M. Stell (2010). Interneurons of the cerebellar cortex toggle Purkinje cells between up and down states. Proceedings of the National Academy of Sciences 107(29), 13153–13158.

29. Person, A. L. and I. M. Raman (2012a). Purkinje neuron synchrony elicits time-locked spiking in the cerebellar nuclei. Nature 481 (7382), 502–505.

30. Person, A. L. and I. M. Raman (2012b). Synchrony and neural coding in cerebellar circuits. Frontiers in neural circuits 6, 97.

31. Schonewille, M., S. Khosrovani, B. H. J. Winkelman, F. E. Hoebeek, M. T. G. De Jeu, I. M. Larsen, J. Van der Burg, M. T. Schmolesky, M. A. Frens, and C. I. De Zeeuw (2006). Purkinje cells in awake behaving animals operate at the upstate membrane potential. Nature Neuroscience 9(4), 459–461; author reply 461.

32. Sugihara, I., H. Fujita, J. Na, P. N. Quy, B.-Y. Li, and D. Ikeda (2009). Projection of reconstructed single purkinje cell axons in relation to the cortical and nuclear aldolase C compartments of the rat cerebellum. The Journal of Comparative Neurology 512(2), 282–304.

33. Tank, D. W., M. Sugimori, J. A. Connor, and R. R. Llinas (1988). Spatially resolved calcium dynamics of mammalian purkinje cells in cerebellar slice. Science 242 (4879), 773â777. PMID: 2847315.

34. Yartsev, M. M., R. Givon-Mayo, M. Maller, and O. Donchin (2009). Pausing Purkinje cells in the cerebellum of the awake cat. Frontiers in Systems Neuroscience 3, 2.

